# Changing feeding levels reveal plasticity in elasmobranch life history strategies

**DOI:** 10.1101/2024.07.11.601909

**Authors:** Sol Lucas, Per Berggren, Ellen Barrowclift, Isabel M. Smallegange

## Abstract

Life history strategies are shaped by phylogeny, environmental conditions and individual energy budgets, and have implications for conservation biology. We summarised life history traits of 151 elasmobranch species into life history strategies for two contrasting feeding levels in a principal components analysis. Two axes, reproductive output and generation turnover, structure elasmobranch life history strategies, contrasting with results from similar studies. Species’ positions in this life history space were not fixed, but shifted to higher reproductive output when feeding level increased. We also found that both axes predicted population performance, but that population growth rate does not necessarily inform on a species’ demographic resilience. Finally, neither axis predicted IUCN conservation status. Our analyses reveal plasticity in species life history strategies and warn against extrapolating the life history strategy framework from one environment to another when predicting a species’ response to (climate) change, perturbations, and (over)exploitation.

## 1. Introduction

Life history strategies of many animals and plants reflect trade-offs at the species level. For example, the fast-slow continuum represents the trade-off between reproduction versus survival (Stearns, 1983), often irrespective of body size or phylogenetic relatedness (Williams, 1966; Gadgil et al., 1970; Reznick, 1983; Gaillard et al., 2016). Yet, trade-offs also operate within individuals, such as trading off growth versus reproduction in the energy budget (Gadgil et al., 1970; Reznick, 1983), as different dynamics can exist between populations or species with similar traits (Nilsen et al., 2009; Gamelon et al., 2021). Specifically, the classical association between life history strategies and population responses to environmental change can break down in variable environments when accounting for individual-level trade-offs and energy allocation during the development of individuals (Rademaker et al., 2024). One reason for this could be that individual-level plasticity allows organisms to adjust energy allocation in response to environmental variability, maintaining stability under resource limitation and accelerating growth and reproduction in favourable conditions (Weidner et al. 2020).

Calls exist for more theory-driven studies of how plasticity in development shapes life history variation (Stott et al. 2024). Additionally, we have limited understanding of how ecological and evolutionary factors, like phylogeny or habitat, shape variation in life history strategies across species (Gaillard et al., 2016; Salguero-Gómez et al., 2016; Salguero-Gómez, 2017; Capdevila et al., 2020a). Identifying patterns and plasticity in life history strategies across the tree of life is one way to predicting population growth rates and demographic resilience (combinedly referred here as ‘population performance’) (Salguero-Gómez et al., 2016). However, to do so requires in-depth understanding of how energy budgets, phylogeny and habitat structure life history strategies across species (Salguero-Gómez, 2017; Capdevila et al., 2020a; Romeijn & Smallegange, 2022; Stott et al. 2024). Elasmobranchs (sharks, skates and rays) encompass a vast amount of life history variation. Longevity, for example, can range from five (e.g. *Carcharhinus sealei* and *Carcharhinus sorrah*; Ebert et al. 2021) to almost 400 years (*Somniosus microcephalus*; Nielsen et al. 2016). Reproduction is highly variable, including oviparity (skates and three families of shark), aplacental viviparity (in various forms; sharks and rays) and placental viviparity (some sharks) (Carrier et al., 2004; Miller et al., 2022). Elasmobranch population performance typically varies with body size, reproductive strategy and habitat (Pardo & Dulvy, 2022; Barrowclift et al., 2023; Gravel et al., 2024), yet little is known about the impact of energy budgets.

Here, we investigate how individual energy budgets, phylogeny and habitat structure elasmobranch life history strategies across environments that differ in food availability, and if the resulting life history framework links to population performance and conservation biology (Salguero-Gómez, 2016; Salguero-Gómez et al., 2017). Using Dynamic Energy Budget Integral Projection Models (DEB-IPMs) for 157 elasmobranch species (Kooijman & Metz, 1984; Sousa et al., 2010; Ellner et al., 2016; Smallegange et al., 2017; Smallegange & Lucas, 2024), we model growth and reproduction based on species-specific energy budgets under different feeding levels (Kooijman, 2001; Kooijman et al., 2008). While regression analyses test predefined trait relationships (Sol et al., 2012; Gaillard et al., 2016; Tomasek et al., 2018), principal component analysis (PCA) identifies emergent life history strategies by summarising trait variation into major axes without prior assumptions (Gaillard et al., 2016; Salguero-Gómez et al., 2016; Paniw et al., 2018; Healy et al., 2019; Capdevila et al., 2020a). Therefore, to identify life history strategies, we (objective 1) calculate, using the parameterised DEB-IPMs, a set of representative life history traits based on schedules of survival, growth and reproduction for a low and a high feeding level (reflecting low and high food availability, respectfully). We (objective 2) summarise these traits using a phylogenetically-corrected PCA to identify major axes of elasmobranch life history strategies. We then (objective 3) assess the influence of phylogenetic ancestry (clade) and habitat (water temperature) on species’ positions along these major axes, and to what extent the position of species along these axes changes with food availability (showing plasticity). Finally, (objective 4) we test whether species position along these axes predicts population growth rate, speed of recovery from perturbations (demographic resilience) and IUCN conservation status. Our approach allows us to investigate whether feeding level influences life history strategies and the resulting conservation biology of elasmobranch species.

## 2. Methods

### 2.1 Brief description of the DEB-IPM

A DEB-IPM describes the dynamics of a population comprising cohorts of females of different lengths that survive, grow and reproduce (Smallegange et al., 2017; Smallegange & Lucas, 2024). A DEB-IPM has eight parameters (Fig. 1: top-left box) and we parameterised DEB-IPMs for 157 elasmobranch species, from 11 orders and 37 families, using the DEBBIES database (Smallegange & Lucas, 2024; with the database downloadable at Smallegange, 2020).

**Figure 1.**
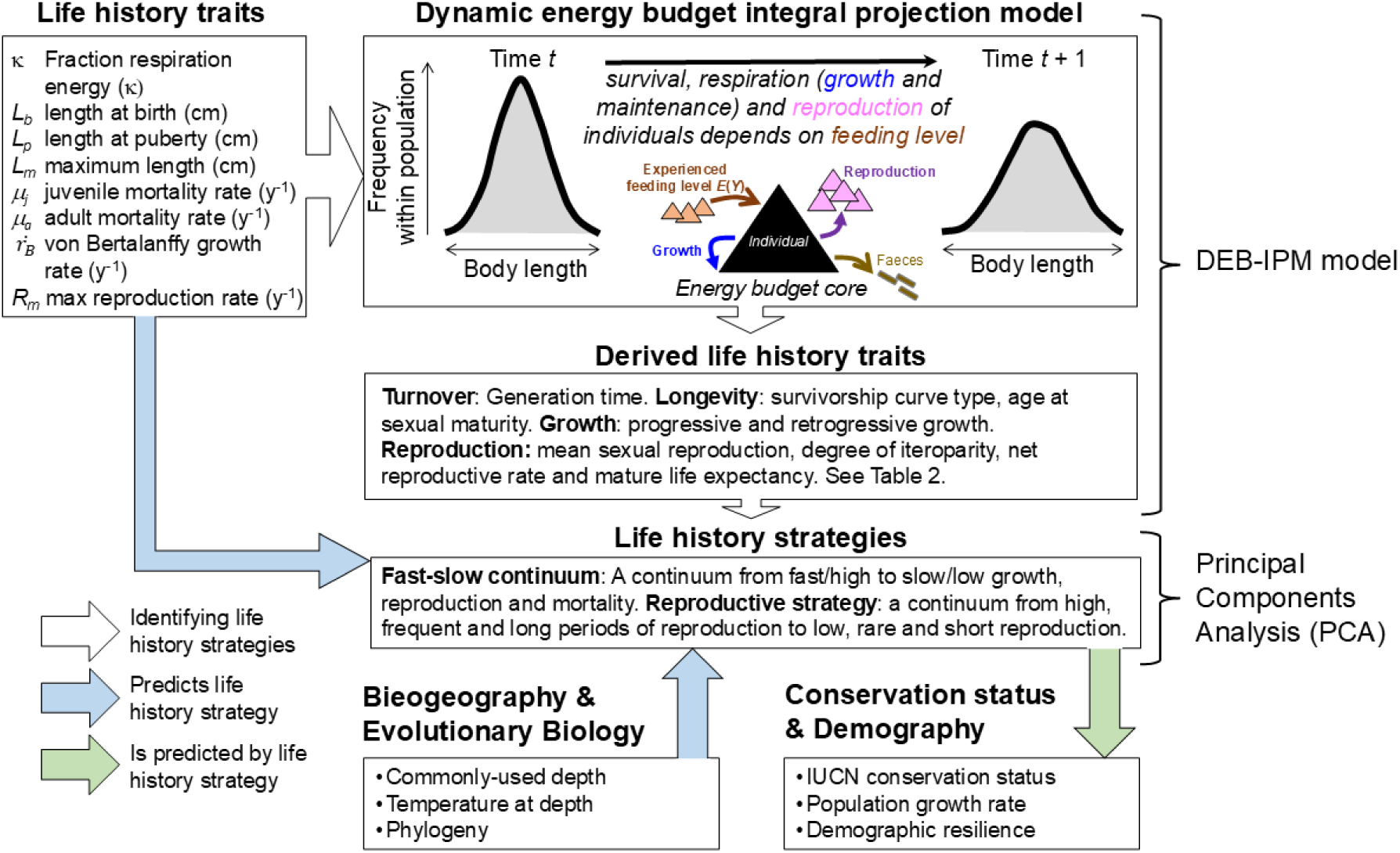
Workflow of parameterising a DEB-IPM using the DEBBIES database (Smallegange & Lucas, 2024), using the derived life history traits to plot elasmobranch life history strategies in a Principal Components Analysis (PCA). Species biogeography and phylogeny were used to estimate position on PCA axes, and the PC scores were used to predict conservation status, population growth rates and demographic resilience. White arrows indicate parameterisation of models or calculation of traits. Blue arrows represent metrics used to predict life history strategies and green arrows indicate the use of life history strategies to make predictions of conservation status and demography. Adapted from Smallegange & Lucas (2024).

In brief, the demographic functions that describe growth and reproduction in a DEB-IPM are derived from the Kooijman-Metz model (Kooijman & Metz, 1984) (see Appendix, section 1 in Figshare repository: Lucas et al., 2024), which is a simple version of the standard model of Kooijman’s DEB theory and only includes the tradeoff between growth and reproduction, but still fulfils the criteria for general explanatory models for the energetics of individuals (Sousa et al., 2010). The Kooijman-Metz model assumes that organisms are isomorphic, are born at length *L*_b_, mature at length *L*_p_ and ingest food at a rate that is proportional to their experienced feeding level *E*(*Y*). This experienced feeding level reflects food availability according to observed gut fullness for fish, ranging from completely empty (0 ≤ *E*(*Y*) < 0.1), to very few food particles (0.1 ≤ *E*(*Y*) < 0.3), to ‘contents divided in different patches’ (0.3 ≤ *E*(*Y*) < 0.7), to ‘just filled (0.7 ≤ *E*(*Y*) < 0.9), to ‘completely full (0.9 ≤ *E*(*Y*) ≤ 1) (Piet & Guruge, 1997). For example, in reef manta rays (*Mobula alfredi*) a feeding level of *E(Y)* = 0.8 is associated with natural, stable populations (Smallegange et al., 2017). At the highest feeding level of *E*(*Y*) = 1, individuals grow to a maximum length *L*_m_. As individuals feed and uptake energy, a constant fraction κ of assimilated energy is allocated to respiration to cover maintenance costs and somatic growth. The remaining fraction 1 – κ of assimilated energy is allocated to reproduction in case of adults and to the development of reproductive organs in case of juveniles. If an individual survives from year *t* to year *t* + 1 (juveniles have a different mortality rate *μ*_j_ than adults, which mortality rate equals *μ*_a_) it grows in length following a von Bertalanffy growth curve characterized by *ṙ*_*B*_; the von Bertalanffy growth rate. If a surviving female is an adult, she also produces offspring at a maximum rate of *R*_m_ if she experiences the highest feeding level of *E*(*Y*) = 1. Note that *R_m_* is proportional to (1 – *κ*), whereas *L_m_* is proportional to κ, which controls energy conservation. However, the role of κ in a DEB-IPM is mostly implicit, as κ is used as input parameter only in the starvation condition (see Appendix, section 1 in Figshare repository: Lucas et al. 2024), whereas *R_m_* and *L_m_* are measured directly from data.

### 2.2 Life history strategies

Eight representative life history traits were calculated based on schedules of survival, growth and reproduction (Table 1) (Salguero-Gómez et al., 2016; Salguero-Gómez, 2017; Capdevila et al., 2020a) in MATLAB Version 9.12 (The MathWorks Inc., 2022) using the code in Smallegange & Lucas 2024 (objective 1). For each species, this was done for a low and high feeding level, *E(Y)*= 0.6 and *E(Y)*= 0.9, reflecting contrasting levels of food availability. *E*(*Y*) = 0.6 reflects unfavourable feeding conditions, whereas a feeding level of *E*(*Y*) = 0.9 represents a full gut and thus favourable food conditions (Piet & Gurunge, 1997). Feeding levels lower than *E*(*Y*)= 0.6 were excluded because these were too low for individual growth and reproduction to occur for many species. Specifically, at the length where *L* > *L*_*m*_*E*(*Y*), individuals shrink and might not reach their maturation size (*L*_p_). Furthermore, individuals die of starvation at length *L* > *L*_*m*_*E*(*Y*)/*κ* (Appendix, section 1 in Figshare repository: Lucas et al. 2024). If *E*(*Y*) = κ, then individuals die of starvation at their maximum length. For most species, we set κ = 0.8 (Smallegange, 2020). Thus, for *E*(*Y*) values (much) lower than 0.8, individuals (much) smaller than the maximum length, die of starvation. These factors combined mean that as feeding level drops, more species experience declining population growth rate and extinction.

**Table 1.** Demographic quantities and their loadings onto the Principal Components axes. To calculate life history traits, we discretised each DEB-IPM (Eqn 1) by dividing the length domain Ω into 200 very small-width discrete bins, resulting in a matrix **A** of size *m* × *n*, where *m*= *n*= 200, and which dominant eigenvalue equals *λ*. Mean lifetime reproductive success *R*_0_ is the dominant eigenvalue of the matrix **F** = **V**(**I** − **GS**)^−1^, where **I** is the identity matrix and **V**= **DR**, with **D** as the parent-offspring association, **R** the reproduction, **G** the growth and **S** the survival matrix (Caswell, 2001); this gives generation time *T*= log(*R*_0_)/log(*λ*) (Caswell, 2001). The mean life expectancy, *η*_e_, is calculated as *η*_e_= 1^T^**N**, where 1 is a vector of ones of length *m* and **N** is the fundamental matrix **N** = (**I** − **S**)^−1^. The longevity of an individual of length *L* is *η*_L_, which means we can calculate age at sexual maturity ***L***_***α***_ = ***η***_***L***_***b***__ and mature life expectancy ***L***_𝝕_ = ***η***_***L***_***p***__ so that ***η***_***e***_ = ***L***_***α***_ + ***L***_𝝕_ (Caswell, 2019, eqn 4.21). *l*_x_ is the probability of surviving to age at least *x*, and *m*_x_ is the average fertility of age class *x* (cf. Tuljapurkar et al., 2009), **G̅** is the mean of G, **V̅** is the mean of V, and *i* and *j* are the row and column entries of the matrix, respectively. The vital rates included in the studied set of demographic quantities (progressive growth *γ*, retrogressive growth *ρ*, and sexual reproduction *φ*) were averaged across the columns *j* (the length bins), weighted by the relative contributions of each stage at stationary equilibrium. For example, to calculate mean sexual reproduction *φ*, we summed the values in the columns *j* of the **V** matrix and multiplied each *φ_ij_* by the corresponding *j*th element *w*_j_ of the stable stage distribution *w*, calculated as the right eigenvector of **A**. Only loadings for the pPCA and mass-corrected PCA are reported, as Pagel’s λ >0.25. Kaiser criterion >1 only for PC1 and PC2, so PC3s were not retained.

| PCA type<br>Number of populations in analysis |  |  |  | pPCA<br>214 (151 species) |  |  | Mass-corrected pPCA<br>168 (117 species) |  |  |
| --- | --- | --- | --- | --- | --- | --- | --- | --- | --- |
| Demographic quantity | Symbol | Definition | Equation | p-PC1 | p-PC2 | p-PC3 | p-PC1 | p-PC2 | p-PC3 |
| Generation time | $T$ | Number of years required for the individuals of a population to be fully replaced by new ones | $T = \frac{\log(R_0)}{\log(\lambda)}$ | **0.912** | -0.149 | 0.295 | **0.517** | **-0.774** | 0.300 |
| Age at maturity | $L_\alpha$ | Number of years that it takes an average individual in the population to become reproductive | $L_\alpha = \eta_{L_b}$ | **0.814** | 0.353 | 0.048 | -0.064 | -0.203 | 0.949 |
| Progressive growth | $\gamma$ | Mean probability of growing to a larger length across the length domain $\Omega$. | $\gamma = \sum_i^m \bar{G}_{i,j} \vert i < j$ | **0.877** | 0.039 | 0.029 | -0.083 | **-0.945** | 0.234 |
| Retrogressive growth | $\rho$ | Mean probability of growing to a smaller length across the length domain $\Omega$. | $\rho = \sum_i^m \bar{G}_{i,j} \vert i > j$ | -0.128 | **-0.890** | 0.196 | **0.889** | 0.173 | -0.223 |
| Mean recruitment success | $\varphi$ | Mean per-capita number of recruits across the length domain $\Omega$. | $\varphi = \sum_i^m \bar{V}_{i,j}$ | **-0.573** | **0.699** | -0.334 | **-0.921** | 0.283 | -0.008 |
| Degree of iteroparity | $S$ | Coefficient of variation in age at reproduction | $S = \bar{m}_x / \sigma(m_x)$ | -0.068 | 0.250 | -0.963 | **-0.888** | 0.062 | 0.086 |
| Net reproductive rate | $R_0$ | Mean number of recruits produced during the mean life expectancy of an individual in the population. | $R_0 = \sum_{x=0}^{x=\eta_e} l_x m_x$ | 0.288 | **0.936** | -0.096 | **-0.797** | -0.054 | 0.546 |
| Mature life expectancy | $L_\omega$ | Number of years from the mean age at maturity ($L_\alpha$) until the mean life expectancy ($\eta_e$) of an individual in the population. | $L_\omega = \eta_{L_p}$ | **0.914** | 0.208 | -0.135 | -0.312 | -0.464 | 0.754 |
| Axis |  |  |  | **Turnover** | **Repro** | - | **Repro** | **Turnover** | - |
| Proportion of variance explained |  |  |  | 45.5% | 34.8% | 8.8% | 48.3% | 34.9% | 7.8% |
| Cumulative proportion of variance explained |  |  |  | 45.5% | 80.3% | 89.1% | 48.3% | 83.2% | 91.0% |
| Kaiser criterion |  |  |  | 13.28 | 7.76 | 0.50 | 14.9 | 7.79 | 0.39 |
| Pagel's lambda (for the pPCA) |  |  |  |  | 0.417 |  |  | 0.335 |  |

To identify life history strategies along major axes (objective 2), we performed a varimax-rotated, phylogenetically corrected principal component analysis (pPCA) (Revell, 2012), using the *phy.pca* function in the R library *phytools* (Revell, 2012; Chamberlain & Szöcs, 2013; Salguero-Gómez et al., 2016; Paniw et al., 2018; R Core Team, 2023). The phylogenetic correction was conducted at the species level, where for each of the 63 species that were present in both the low and high feeding levels, each duplicate was treated as a separate population of the same species using a dichotomous branch at the species’ tip (Paniw et al., 2018; Healy et al., 2019). Traits were log-transformed and scaled to adhere to PCA assumptions (μ= 0 and SD= 1).

To correct for phylogenetic relatedness among species, a pool of 10,000 possible phylogenetic trees were obtained from Stein et al. (2018), available at Vertlife.org. These trees represented 500 root node ages combined with 20 scenarios for infilling taxon with no genetic data. We randomly selected twenty trees (Barrowclift et al., 2023) and ran the pPCA for each. Each rooted phylogenetic tree had its branch lengths scaled proportionally based on time of separation of clades and species, using the *rotl* package (Michonneau et al., 2016). The pPCA links the phylogeny to the life-history traits via a modified covariance matrix and estimated Pagel’s *λ*, a scaling parameter for the phylogenetic correlation between species, expected under Brownian motion (Freckleton et al., 2002). Pagel’s *λ* was estimated using the *ape* package (Paradis & Schliep, 2019) and varies between 0 (phylogenetic independence) and 1 (species traits covary proportionally to their shared evolutionary history) (Revell, 2010). The loadings for each axis were consistent across all trees (see Appendix, File 1 in Figshare repository: Lucas et al. 2024) and Pagel’s *λ* was consistently above *λ*>0.25 (see Appendix, File S2 in Figshare repository: Lucas et al. 2024), which we set as our cut-off value, above which we assumed the phylogenetic signal indicates meaningful evolutionary relationships. Therefore, we report results from one single tree, randomly selected from the subsample of twenty trees (see Appendix, Fig S1 in Figshare repository: Lucas et al. 2024). Six species were excluded from the analyses as they were either not in the Vertlife.org database (*Squalus hawaiiensis*, *Aetobatus narutobiei* and *Maculabatis ambigua*), they did not have an associated polygon shapefile (see below) on the IUCN database (*Etmopterus granulosus* and *Aetomylaeus bovinus*), or were identified as an extreme outlier after visual inspection of the PCA due to extreme life history trait values compared to other species in the dataset (*Somniosus microcephalus*).

We checked the influence of phylogeny and body mass on the structuring of life history strategies (see Jeschke & Kokko 2009 for a detailed discussion) by running PCA analyses with and without a phylogenetic correction, and with and without a body mass correction. Body mass was corrected for by computing the residuals for linear models between the log_10_-transformed life history traits (Table 1) and body mass for each species for both the PCA and the pPCA (Jolicoeur et al., 1984; Gaillard et al., 1989; Revell, 2009). Body mass values were extracted from Fishbase (Froese & Pauly, 2023) using the *rfishbase* package (Boettiger et al., 2012). Where available, we used the maximum mass given on Fishbase. Otherwise, we employed length-weight relationships to infer maximum body mass. If the length-weight relationship used a length type that was not in the database, length-length conversions were conducted prior to length-weight conversions. Data were not available for 34 species, so there were 151 species in the main analyses and 117 species in the analyses containing body mass corrections (Table 1). We set the same cutoff value of 0.25 for Pagel’s *λ*, above which we used the pPCA, followed by a qualitative check of loadings between the pPCA and the mass-corrected pPCA. Where the mass-corrected loadings were qualitatively different from loadings of analyses without the mass-correction (i.e. where traits were loaded onto different axes), we continued with the mass-corrected analyses. We assessed the significance of PC axes using Kaiser’s criterion, retaining PC axes with eigenvalues greater than unity (Kaiser, 1960).

To compare if and how species position in the life history space changed between the low and high feeding levels, we conducted Pearson’s product-moment correlation tests, between feeding level (*E(Y)*) and PC scores for each axis.

### 2.3 Effect of clade and habitat (water temperature) on life history patterns

We assessed if habitat and clade affect elasmobranch life history variation (objective 3). We sourced habitat information (water depth and temperature) for each species. Continuous variable ‘commonly used depth’ was sourced for each species from the Fishbase database (Boettiger et al., 2012; Froese & Pauly, 2023). If commonly used depth was unavailable, median depth was sourced from the IUCN database (IUCN, 2021). For temperature, we used ‘temperature-at-depth’ following the methodology of (Barrowclift et al., 2023). To this end, shape files containing polygons of known depth ranges for each species were taken from the IUCN database (IUCN, 2021). These were overlaid onto the International Pacific Research Center’s mean annual ocean temperatures across 27 depth levels (0-2000m), based on a dataset from the Argo Project (available at http://apdrc.soest.hawaii.edu/projects/Argo/data/statistics/On_standard_levels/Ensemble_mean/1x1/m00/index.html). Median temperature-at-depth was calculated using values at the nearest depth level to each species’ commonly used depth (or the IUCN median depth where commonly used depth was unavailable). For regionally endothermic species (family *Lamnidae,* Dickson & Graham, 2004) a correction factor of 3.5°C was applied to their median temperature, consistent with a similar demographic study by Pardo and Dulvy (2022).

We tested for collinearity between depth, temperature and clade using visual inflation factor (threshold value VIF>10) in the *faraway* R package (Faraway, 2022). Depth, temperature and clade were all colinear. Therefore, we separately assessed the effect of temperature at depth (as the representative variable for habitat) and clade on the PC scores of the first and second axis of the (p)PCA. We used two general linear mixed-effect models (LMERs) where the predictor variable was either temperature at depth or clade, with a random effect of species in each model, and the response variable was either PC1 or PC2 score. For each LMER, the model assumptions of Gaussian errors and homoscedasticity were confirmed by inspecting the probability plots and error structures in R (Lüdecke et al., 2021; Faraway, 2022; R Core Team, 2023).

### 2.4 Life history strategies as predictors of population growth rate, demographic resilience and conservation status

We used linear models (LMs) to test if the PC scores (continuous variables) of the first and second axis of the best fitting PCA, and their interaction, predict population growth rate (*λ*) and demographic resilience across species (objective 4). Even though PCA assumes that the principal components (PC1 and PC2) are uncorrelated, this assumption pertains to the original data’s variance structure. By testing interactions between PC1 and PC2 we explored whether the combined influence of these principal components provides additional explanatory power for our response variable, beyond their individual effects. To calculate *λ*, we discretised each DEB-IPM (Eqn 1) by dividing the length domain Ω into 200 very small-width discrete bins, resulting in a matrix **A** of size *m*×*n*, where *m*= *n*= 200, and which dominant eigenvalue equals *λ* (Table 1). Demographic resilience was calculated as the damping ratio *ξ*, with *ξ* = *λ*/||*λ*_2_||, where *λ*_2_ is the highest subdominant eigenvalue of matrix **A** (Caswell, 2001; Capdevila et al., 2020b). Alternative metrics exist to capture resilience (simulated time-to-recovery or transient growth rates [Capdevila et al. 2021]) but for ease of comparison with previous studies we used the damping ratio. To investigate predictive links between life history strategies and conservation status, global conservation statuses for each species were sourced from the IUCN database (IUCN, 2021), where LC indicates Least Concern, NT indicates Near Threatened, VU indicates Vulnerable, EN indicates Endangered, CR indicates Critically Endangered and DD indicates Data Deficient. We used an ordinal regression to assess if PC scores predict species IUCN conservation status (excluding Data Deficient species).

All LMs, LMERs, ordinal regression analyses and plots (Wickham, 2016; Attali & Baker, 2023; Hijmans, 2023; Slowikowski, 2023) were performed in R version 4.3.2 (R Core Team, 2023) in Rstudio (RStudio Team, 2023), using the *dplyr* and *tibble* packages for data manipulation (Wickham et al., 2022; Müller & Wickham, 2023). For each LM, the model assumptions of Gaussian errors, homoscedasticity and collinearity were confirmed by inspecting the probability plots and error structures in R (Lüdecke et al., 2021; Faraway, 2022; R Core Team, 2023).

## 3 Results

### 3.1 Two axes structure elasmobranch life history strategies (objective 1 & 2)

We found that two separate PC axes structure elasmobranch life history traits, cumulatively explaining 83.2% of the total variance in life histories (Table 1: PC1: 48.3%, PC2: 34.9%). Pagel’s *λ* was higher than our cut-off value of 0.25 (Pagel’s *λ*= 0.31), indicating the phylogenetic signal was significant. In addition, correcting for body mass changed the dominant axes (Table 1) and we thus continued with the mass-corrected pPCA. Degree of iteroparity (*S*), mean recruitment success (*φ*) and net reproductive rate (*R_0_*) were loaded negatively onto PC1, whereas retrogressive growth (*ρ*) and generation time (T) were loaded positively (Fig. 2A) (Table 1). These life history traits are all associated with reproductive output. Thus, as PC1 scores move from positive to negative, elasmobranchs attain greater lifetime reproductive success while their retrogressive growth and generation times decrease (Fig. 2A). Generation time (*T*) and progressive growth (*γ*) were loaded negatively onto PC2 (Fig. 2A) (Table 1). As PC2 scores move from positive to negative, individuals increase their allocation to growth while population turnover decreases (i.e., greater generation time), so this axis is described as ‘generation turnover’ (Fig. 2A). Finally, mean age at maturity (*L_α_*) and mature life expectancy (*L_ω_*) were not loaded onto either PC1 or PC2 (Table 1). The Kaiser criterion did not meet the threshold (<1) for PC3, so this axis did not explain sufficient variance in life history strategies to be included in further analyses.

**Figure 2.**
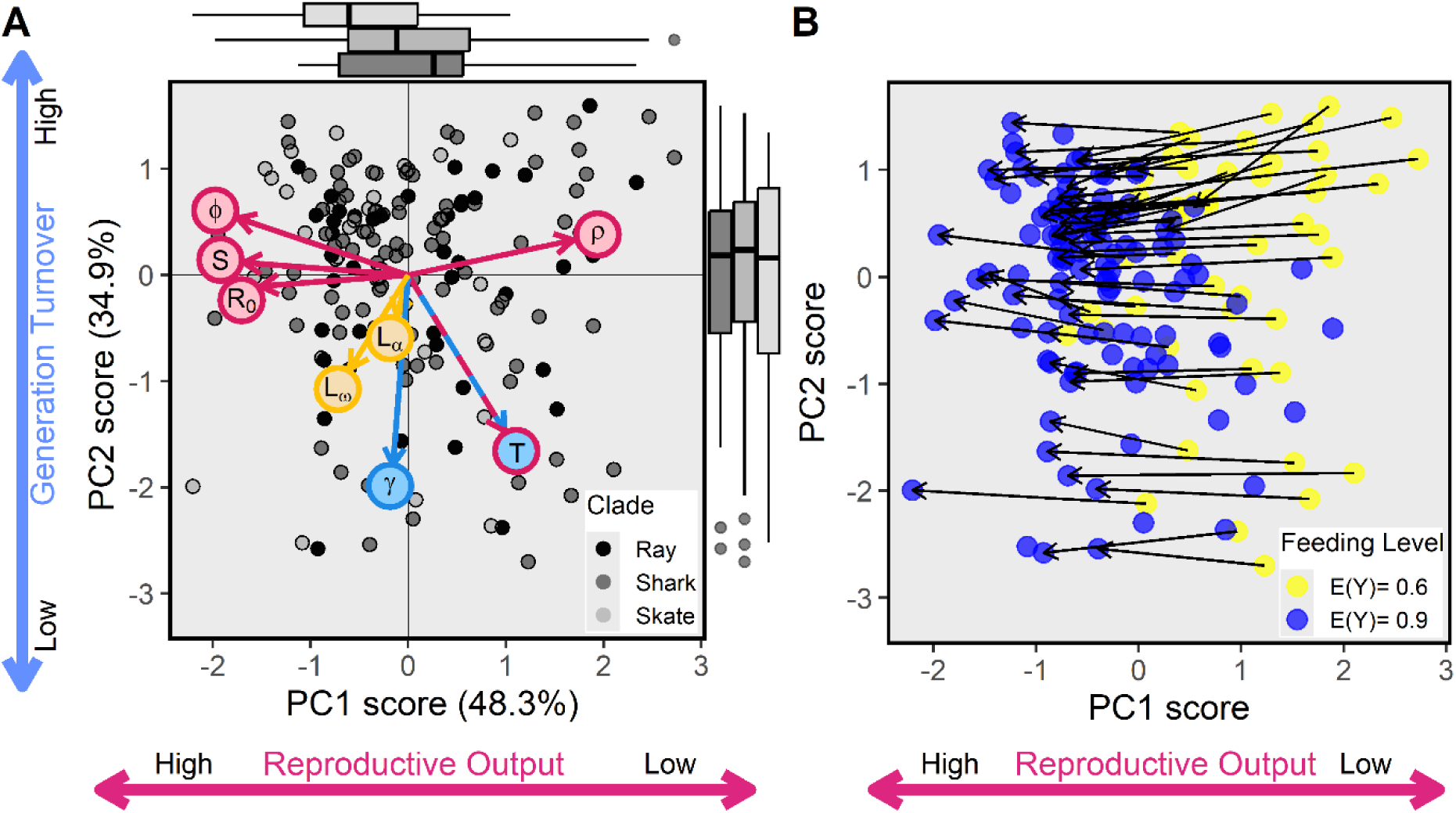
Mass-corrected (p)PCA plots including 117 species from the high feeding level model (E(Y) = 0.9) and 51 of these species from the low feeding level model (E(Y) = 0.6). The 51 species that were in both high and low feeding level models were treated as independent populations, leading to a total of 168 data points in each plot (A) and (B). (A) Marginal boxplots of the split by clade (Shark, Skate and Ray) are included for each principal component. Labelled arrows correspond to the variables used in the PCA analysis and show the magnitude and direction of the loadings. Magenta arrows align with the reproductive axis, blue arrows align with the generation turnover axis and yellow arrows are not aligned with either PC1 or PC2. The dashed blue/magenta arrow aligns with both the reproductive output and generation turnover axes. (B) The change in PC scores is marked by arrows for 51 species in both low (yellow) and high (blue points) feeding level. 66 species appear only in blue, with no connected arrow. The change in life history strategies between low and high feeding level creates a shift towards higher reproductive output.

Feeding level was negatively correlated with PC1 score (reproductive output), r(166) = -0.639, p = <0.001. As feeding level increases, PC1 scores decreased, corresponding to an increase in lifetime reproductive output between the low and high feeding level as species can spend more energy to reproduce at a higher rate (Fig 2B). There was no correlation between PC2 scores (generation turnover axis) and feeding level (p = 0.186).

### 3.2 Habitat type shapes life history strategies differently for different feeding levels and clades (objective 3)

Temperature did not significantly affect PC1 score (reproductive output) (p= 0.423). There was a significant effect of Clade on PC1 score (F_2,165_= 4.12, Marginal R^2^= 0.047, p= 0.008). PC1 scores were lower for skates than rays and sharks, indicating that skates have a high reproductive output (Fig. 2A). PC2 scores (generation turnover) increased with increasing temperature at depth (F_1,116_=, Marginal R^2^= 6.94, p= 0.010: Fig. 3B). This suggests that across all elasmobranchs, species with faster life history strategies occur in warmer waters. Phylogenetic clade did not significantly affect PC2 score (p = 0.762).

**Figure 3.**
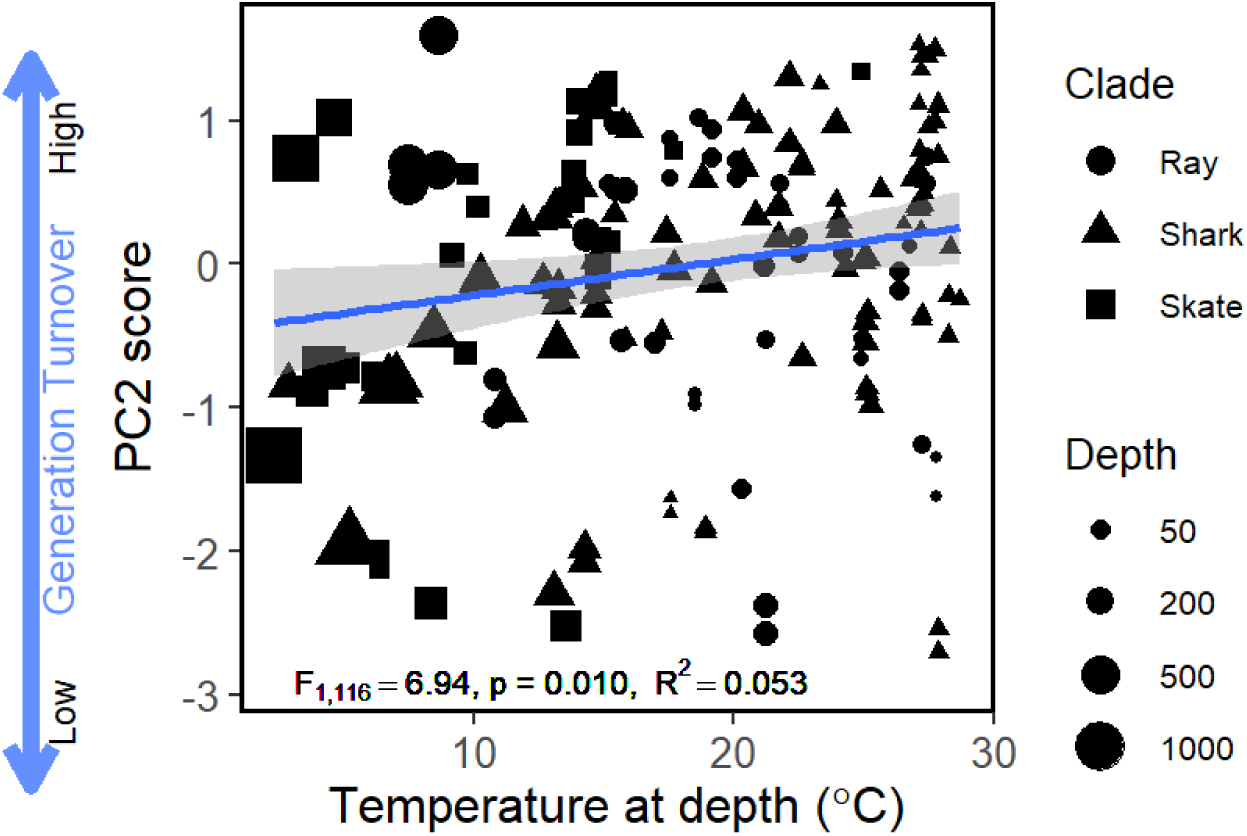
PC2 score decreased for increasing temperature-at-depth. Warmer temperatures relate to positive PC2 scores (higher generation turnover). The F-statistic, p-value and Marginal R^2^ (variance explained by fixed effects only) are in the plot window. Conditional R^2^ (variance explained by fixed and random effects) was 0.963. The grey band around the fitted models shows the 95% confidence interval.

### 3.3 Life history strategies predict population performance, but not IUCN status (objective 4)

Population growth rate was significantly affected by the interaction between PC1 scores (reproductive output) and PC2 scores (generation turnover) (Table 2). Population growth rate increased with decreasing PC1 scores, but this increase was steeper for species with high, positive PC2 scores (higher turnover) than for species with high, negative PC2 scores (lower turnover) (Fig. 4A). Resilience to perturbations (damping ratio) significantly increased with decreasing PC1 scores (reproductive output) and increasing PC2 scores (generation turnover) (Table 2), showing that species with higher reproductive outputs and higher generation turnover have higher demographic resilience (Table 2, Fig. 4B). There was no significant association of PC1 scores, PC2 scores, or their interaction on the IUCN category of species (Table 2, Fig. 4C).

**Figure 4.**
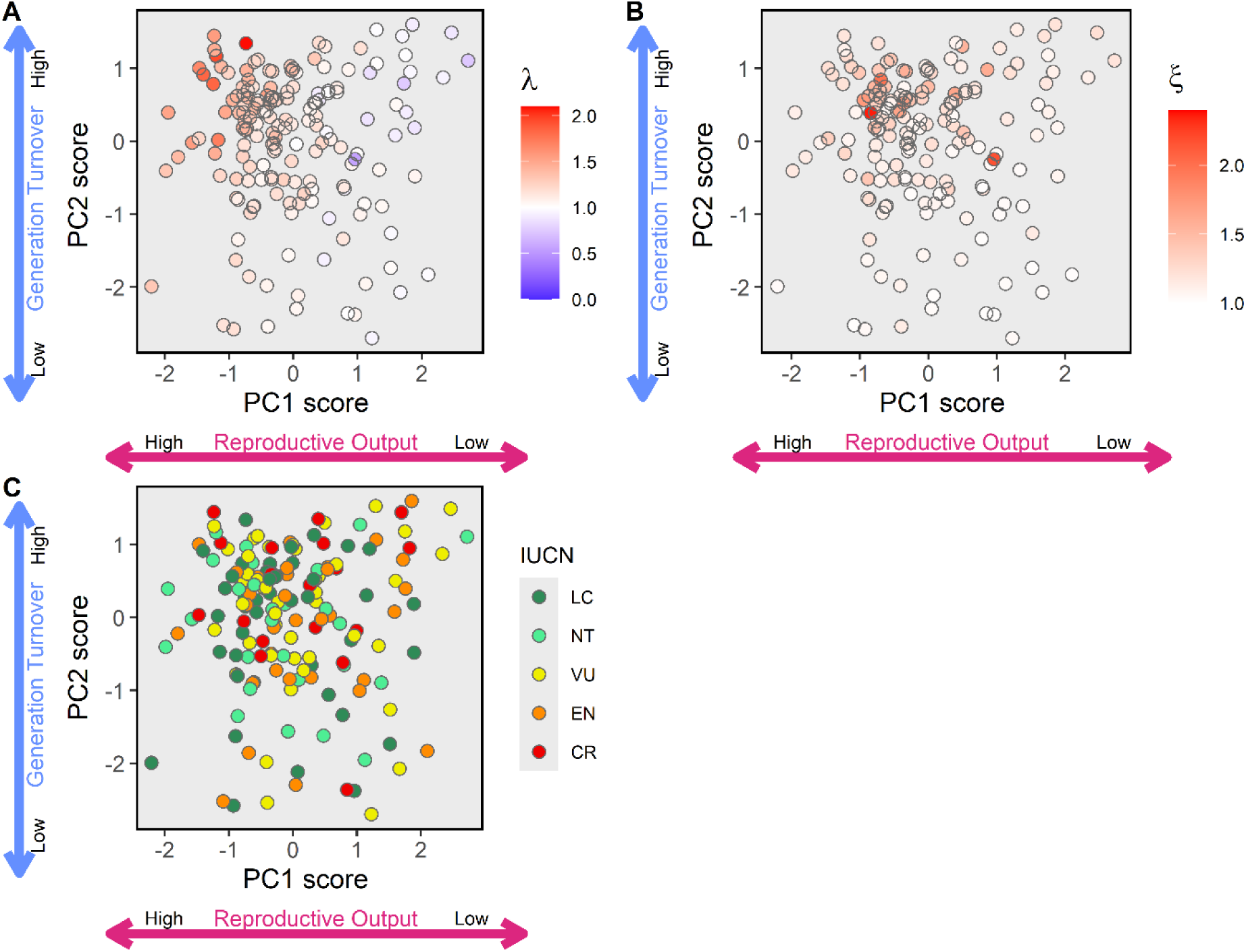
Overlays on the PCA figures for: population growth rate (A), damping ratio (B), IUCN conservation status (C).

**Table 2.**
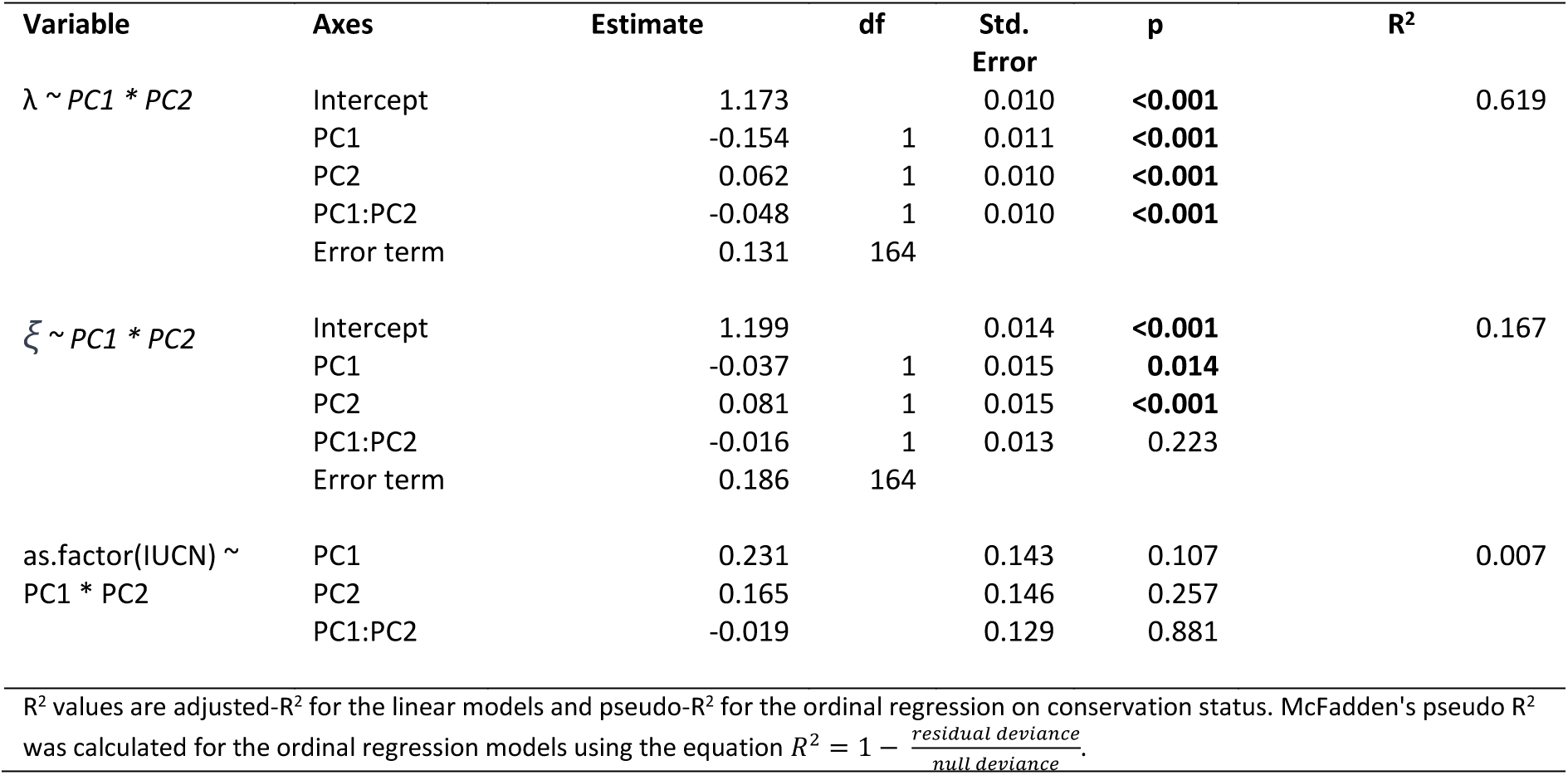
General linear models for population growth rate (*λ*) and demographic resilience (damping ratio, *ξ*), as well as ordinal regression models for IUCN conservation status, as a response to PC1 (reproductive output) and PC2 (turnover) scores, for both the low (*E(Y)*=0.6) and high (*E(Y)*=0.9) feeding levels.

| Variable | Axes | Estimate | df | Std.<br>Error | p | R<sup>2</sup> |
| --- | --- | --- | --- | --- | --- | --- |
| $\lambda \sim PC1 * PC2$ | Intercept | 1.173 | | 0.010 | **<0.001** | 0.619 |
|  | PC1 | -0.154 | 1 | 0.011 | **<0.001** |  |
|  | PC2 | 0.062 | 1 | 0.010 | **<0.001** |  |
|  | PC1:PC2 | -0.048 | 1 | 0.010 | **<0.001** |  |
|  | Error term | 0.131 | 164 |  |  |  |
| $\xi \sim PC1 * PC2$ | Intercept | 1.199 | | 0.014 | **<0.001** | 0.167 |
|  | PC1 | -0.037 | 1 | 0.015 | **0.014** |  |
|  | PC2 | 0.081 | 1 | 0.015 | **<0.001** |  |
|  | PC1:PC2 | -0.016 | 1 | 0.013 | 0.223 |  |
|  | Error term | 0.186 | 164 |  |  |  |
| as.factor(IUCN) ~<br>PC1 * PC2 | PC1 | 0.231 |  | 0.143 | 0.107 | 0.007 |
|  | PC2 | 0.165 |  | 0.146 | 0.257 |  |
|  | PC1:PC2 | -0.019 |  | 0.129 | 0.881 |  |

R<sup>2</sup> values are adjusted-R<sup>2</sup> for the linear models and pseudo-R<sup>2</sup> for the ordinal regression on conservation status. McFadden's pseudo R<sup>2</sup> was calculated for the ordinal regression models using the equation $R^2 = 1 - \frac{\text{residual deviance}}{\text{null deviance}}$ .

## 4 Discussion

Our aim was to investigate how individual energy budgets, phylogeny and habitat type structure elasmobranch life history variation, and if this structured variation can predict population performance and conservation biology. Elasmobranch life histories are captured by a reproductive output and generation turnover axis. Interestingly, we found that as feeding level increased, species moved towards higher reproductive output along the reproductive output axis but not the generation turnover axis. Regardless, the two life history axes predicted species’ population growth rate and demographic resilience, but not IUCN conservation status.

### 4.1 The impact of feeding level on quantifying and applying life history strategies

For many taxa, including elasmobranchs, their life history strategies are influenced by phylogeny and habitat (Capdevila et al., 2020a) as these impose selection on the life history trajectories of individuals. In this study, we captured the phenomenon that species move from lower to higher reproductive outputs as feeding level increases. In the core energy budget of our model, a fraction of the acquired energy is allocated to growth, while the remainder goes to reproduction (Kooijman & Metz, 1984). This trade-off is implicitly represented in our model through the values of maximum length (*L_m_*) and maximum reproduction rate (*R_m_*). Consequently, we cannot determine the actual value of this trade-off or how it is resolved across species. However, we can hypothesise how different resolutions of the growth-reproduction trade-off might influence species life histories as feeding levels increase. If a high proportion of energy is allocated to growth, an increase in feeding level would result in higher values of life history traits associated with growth (progressive growth *γ* and retrogressive growth *ρ*, respectfully) (Smallegange & Guenther, 2025). Conversely, if a high proportion of energy is allocated to reproduction, an increase in feeding level would enhance life history traits associated with reproduction (mean recruitment success *φ*, degree of iteroparity *S*, net reproductive rate *R*_0_ and mature life expectancy *L*ω) (Smallegange & Guenther, 2025). These four reproductive traits align with our reproductive output axis. Therefore, if all elasmobranchs in our study allocate relatively more energy to reproduction than to growth, this would explain why changes in feeding levels are reflected solely along the reproductive output axis. This strong association with reproductive output may explain why our findings differ from previous studies, which have consistently identified the fast-slow continuum (pace of life axis) as the dominant predictor of life history variation across a range of taxa (Appendix: Table S2; Capdevila et al. 2020a; Healy et al. 2019; Salguero-Gómez 2017; Salguero-Gómez et al. 2016), as shifts in feeding level drive changes along the reproductive output axis in our model.

In reality, species will vary in both energy acquisition and allocation as environmental conditions change. Understanding how these factors interact is key to predicting life history variation (Laskowski et al. 2021). For example, a pace of life axis can emerge if species differ in energy allocation at constant acquisition (feeding level), or if they differ in acquisition but have the same, constant allocation (Smallegange & Guenther 2025). However, individuals can also adjust energy allocation during their lifetime, redistributing it for maintaining functional stability in poor conditions or accelerating growth and reproduction when resources are abundant (Weidner et al. 2020). As a result, the same individuals within a population can show plasticity and change from a slow to fast pace of life, or vice versa, as energy availability varies over time (Del Giudice 2014). While within-population links between life history and behaviour (pace of life syndrome [Dammhahn et al. 2018]) are increasingly documented, their transferability across populations and species remains uncertain. For instance, Careau et al. (2009) observed a positive correlation between age at first reproduction and exploratory behaviour across nineteen muroid species, with subsequent studies confirming similar within-population patterns in house mice (Prabh et al., 2023) and common voles (Eccard et al., 2022). These findings suggest that while plasticity in life history and behaviour is evident within populations, its generalisability across taxa requires further investigation (Smallegange & Guenther 2025).

Plasticity in life history strategies can be adaptive when developing organisms adjust their allocation decisions in response to their local (feeding) environment, following evolved rules that maximise expected fitness in different ecological conditions (McNamara & Houston, 1996; Ozgul et al. 2023). But the consequences of such plasticity and differences in the pace of life and other life history strategies across populations and species can be far-reaching. Like others (Salguero-Gómez, et al., 2016; Salguero-Gómez, 2017), we found that two life history strategy axes predict species population growth and resilience. Perhaps unsurprisingly, both population growth and resilience increased with increasing feeding level along the reproductive output axis. However, a high population growth rate did not always equate to a high demographic resilience. For example, species with high reproductive output and generation turnover show high population growth rates but not always high resilience. In fact, complex environmental interactions and changes in pace of life can influence species extinction risk under climate change (Ozgul et al. 2023). However, our analysis did not find a link between life history strategies and IUCN threat status, likely because the IUCN status is fixed per species. Thus, plastic shifts in life history strategies would not predict threat status unless they impact it (as in Ozgul et al. 2023). All taken together, one should therefore be cautious when using life history strategies to predict how species perform in different environments.

In our analyses, progressive growth was higher for species with lower turnover rates, not higher ones (e.g. Salguero-Gómez 2016; Salguero-Gómez et al. 2017) (see Appendix, Table S2 in Figshare repository: Lucas et al. 2024). We surmise that this is due to high variation in the von Bertalanffy growth rate between species, driving a positive correlation between generation time and progressive growth (cf. van Noordwijk & de Jong 1986). For example, the values of progressive growth and generation time change with different values of von Bertalanffy growth rate and feeding level. The relationship between generation time and progressive growth for the reef manta ray, *Mobula alfredi,* can be plotted for a range of feeding levels and values of von Bertalanffy growth rate (see Appendix, Fig. S2 in Figshare repository: Lucas et al. 2024). For the same von Bertalanffy growth rate value, generation time increases with decreasing feeding level and progressive growth decreases. At the same time, generation time decreases with increasing von Bertalanffy growth rate values. The result is that, at the same feeding level (black solid lines in Appendix, Fig. S2 in Figshare repository: Lucas et al. 2024), progressive growth increases with decreasing values of von Bertalanffy growth rate, and simultaneously, generation time increases. Because our dataset covers a wide range of von Bertalanffy growth rate values, this would explain why we found that the slower a species’ life history speed, the higher its apparent progressive growth. These findings highlight the need for careful interpretation of correlations between life history traits calculated from demographic datasets (our objective 1 and e.g. Salguero-Gómez et al., 2016; Salguero-Gómez, 2017; Paniw et al., 2018; Healy et al., 2019; Capdevila et al., 2020a), and how these are used to predict population characteristics (van Noordwijk & de Jong, 1986).

### 4.2 Consequences for elasmobranch conservation

For many oceanic species, demographic data are sparce or lacking (Bradshaw et al., 2007; Pardo et al., 2016a; Temple et al., 2020). This often means the only measure available to evaluate a species relative risk to perturbations like fishing or environmental change, is an estimate of its maximum intrinsic rate of population increase, *r_max_* (Myers et al., 1997, 1999; Dulvy et al., 2004; Pardo et al., 2016b). Typically, species with higher *r_max_* values are assumed to have higher resilience to perturbations (cf. Cortés 2016; Pardo et al. 2016b, a). The intrinsic rate of population increase, *r*, is related to population growth rate (*λ* = *e*^*r*^). We found that species with the highest population growth rates do not necessarily show the highest resilience. This highlights the importance of exercising caution when using summary metrics such as *r_max_* to predict species’ resilience (cf. Smallegange et al. 2020). Our approach presents a more mechanistic methodology that can be applied to data-sparce elasmobranchs to understand and predict their resilience and conservation status.

There is a broad diversity of diet and feeding strategies in elasmobranchs (Cortés et al., 2008), which potentially influence species-specific life history strategy responses to changes in feeding levels. In this study, we found a consistent response between species to increase reproductive output with increased feeding level. However, when assessing individual species, researchers should consider realistic values and changes in feeding levels based on available literature or empirical studies to estimate population responses. Additionally, global (sea water) warming is linked to poorer hunting performance in sharks (Pistevos et al., 2015). Warming can thus reduce the sharks’ experienced feeding level, population growth rate and ultimately their resilience. Understanding how elasmobranch life histories, their habitats and feeding translate into differences in resilience is crucial to informing management decisions and predicting conservation status. We did not find a relationship between life history axes and IUCN conservation status, possibly because plastic shifts in life history strategies were not linked to IUCN status (as discussed above), or because exploitation was not included in our models. Given that the most immediate threat to sharks, skates and rays is overfishing (Dulvy et al., 2021; Pacoureau et al., 2021; Worm et al., 2024), further study using an energy budget, demographic approach like ours could examine the effects of different fisheries mortalities (either by empirical measures or by derived estimates; Smith et al. 1998) on population performance of threatened or highly fished species.

### 4.3 Conclusion

Studies have shown that population responses to future environmental change and perturbations depend on species-specific life-history strategies (e.g., Ozgul et al. 2023). Further research should explore if variations in environmental conditions fuel plasticity of life history strategies in a range of taxa, and how these impact population performance and responses to change. Our analyses reveal that feeding level can cause plasticity in life history strategies, impacting how the dominant axes can be used to predict population responses to change and perturbations. For elasmobranchs, we provide strong support for the expansion of the classical use of maximum intrinsic rate of population increase, *r_max_*, to incorporating major axes of life history strategies, and highlight how our approach can be used to explore different scenarios of (over)fishing to quantify sustainable levels of exploitation.

## Statement of Authorship

SL: Conceptualisation, Methodology, Formal analysis, Investigation, Validation, Data curation, Writing- Original draft preparation, Writing- Review & Editing, Visualisation; PB: Writing- Reviewing and Editing, Supervision; EB: Methodology, Formal analysis, Writing- Reviewing and Editing; IMS: Conceptualisation, Methodology, Formal analysis, Investigation, Validation, Data curation, Writing- Original draft preparation, Writing- Review & Editing, Visualisation, Supervision.

## Data accessibility statement

**Supplementary information**, including data and analysis code used to run the models in MATLAB and RStudio, are available at Figshare (DOI: https://doi.org/10.6084/m9.figshare.26166427). A publicly accessible pre-print for this article is available at: https://doi.org/10.1101/2024.07.11.601909.

## Acknowledgements

SL was funded by the Leverhulme Doctoral Scholarship in Behaviour Informatics (DS-2017-015).

